# Antisense oligonucleotides targeting exon 11 are able to partially rescue the Neurofibromatosis Type 2 phenotype *in vitro*

**DOI:** 10.1101/2022.02.11.479859

**Authors:** N. Catasús, I. Rosas, S. Bonache, A. Negro, M. Torres-Martin, A. Plana, H. Salvador, E. Serra, I. Blanco, E. Castellanos, the NF2 Spanish National Reference Centre HUGTP-ICO-IGTP

## Abstract

Neurofibromatosis type 2 (NF2) is an autosomal dominant condition caused by loss of function variants in the *NF2* gene, which codes for the protein Merlin, and characterized by the development of multiple tumours of the nervous system. The clinical presentation of the disease is variable and related to the type of the inherited germline variant. Here, we tested if PMOs could be used to correct the splice signalling caused by variants at +/-13 within the intron-exon boundary region. Here we show that the PMOs designed for these variants do not constitute a therapeutic approach. Furthermore, we evaluated the use of phosphorodiamidate morpholino oligomers (PMOs) to reduce the severity of the effects of *NF2* truncating variants with the aim of generating milder hypomorphic isoforms *in vitro* through the induction of the in-frame deletion of the exon-carrying variant. We were able to specifically induce the skipping of exons 4, 8 and 11 maintaining the *NF2* gene reading frame at cDNA level. Only the skipping of exon 11 produced a hypomorphic Merlin (Merlin-e11), able to partially rescue the observed phenotype in primary fibroblast cultures from NF2 patients, being encouraging for the treatment of patients harbouring truncating variants located in exon 11.

## INTRODUCTION

Neurofibromatosis type 2 (NF2) (OMIM 101000) is an autosomal dominant (AD) disease caused by loss-of-function (LOF) variants in the *NF2* tumour suppressor gene (22q12.2) ^1,2^. The reported incidence is between 1 in 28,000-40,000^3^ and the hallmark of the disease is the presence of bilateral vestibular schwannomas (BVS) which clinically present with hearing loss, tinnitus and imbalance^4^. Common features of NF2 include cranial, spinal, peripheral nerve, and intradermal schwannomas; cranial and spinal meningiomas; and intrinsic central nervous system (CNS) tumours, usually spinal ependymomas^5,6^. Despite their benign nature, tumours are associated with high morbidity due to their multiplicity and anatomical location, which cause pain and nerve dysfunction leading to a reduced quality of life and life expectancy^4^.

There are limited therapies available for these patients and radiotherapy and surgery for tumour resection are the classical treatments^4^. Currently, medical therapy is also an option since Bevacizumab, a monoclonal antibody targeting the Vascular Endothelial Growth Factor (VEGF), has been demonstrated to be an effective treatment for vestibular schwannomas (VS) inducing the reduction of tumour growth and an improvement in hearing in adult NF2 patients^6,7^. However, Bevacizumab does not seem to be equally effective in all NF2 patients and, in addition, it has no effect on other types of tumours arising from the disease, such as meningiomas^8^. Efforts have been made to develop other therapies for NF2-related tumours, such as Losartan (antihypertensive), Brigatinib and Lapatinib (tyrosine kinase inhibitors) or AR-42 (Histone deacetylase (HDAC) inhibitor)^9–13^, but there still remains a need to identify clinically relevant therapeutic targets to treat NF2 patients.

The *NF2* gene consists of 17 exons^14,15^ with a wide mutational spectrum. It is estimated that 85-90% of the pathogenic variants in *NF2* are germline point mutations and 10-15% are large deletions, distributed throughout the entire gene^16,17^. *NF2* encodes for Merlin (Moesin-Ezrin-Radixin-Like protein), a 595aa scaffold protein involved in membrane protein organization, regulation of cell-cell adhesion and with a regulatory role in the cytoskeleton architecture^18,19^. Merlin is comprised of three structural regions: a N-terminal FERM domain which is subdivided into three globular domains (F1, F2 and F3), an α-helical coiled-coil domain and a C-terminal hydrophilic tail^20^.

The genotype-phenotype correlation of NF2 has long been studied^21–29^. For NF2, it is well reported that truncating variants and large deletions, which account for around 30% of cases, are associated with severe manifestations of the disease, whereas missense or in-frame variants present milder forms of NF2. Data reveal that individuals with the latter variants show a late disease onset and are diagnosed later compared to individuals with truncating mutations, as well as, a reduced risk of developing peripheral nerve tumours, spinal tumours and meningiomas. Thus, being these genetics variants associated with milder disease burden^21,23,27,30,31^. Besides, about 25% of the point mutations affect the correct splicing of the *NF2* gene, and this mechanism is associated with a variable severity of the disease^26,28,30^. NF2 is also estimated to have a high presence of mosaicism, reported to affect at least 30% and up to 50% of *de novo* patients, which often challenges the molecular diagnosis of the disease^32^ NF2 mosaicism is associated to mild clinical presentations, although significant variability has been reported^28,32^. Thus, the well-known association between the germline genetic variant type and disease severity may constitute an opportunity to develop personalized gene therapies based on the specific variant of the *NF2* gene.

Antisense gene therapy has been successfully used to regulate gene expression in some pathologies. Most of the target diseases are inherited in an autosomal recessive manner, such as Spinal Muscular Atrophy (SMA)^33^, homozygous familial hypercholesterolemia (HoFH)^34^, or Duchenne muscular dystrophy (DMD)^35^ the latter two have been approved by the FDA and are amongst others currently in the advanced stages of clinical trials^36,37^. Furthermore, there are also examples for AD diseases, such as Retinitis pigmentosa (RP)^38^, amyotrophic lateral sclerosis^39^ or Huntington’s disease^40^. In all cases, antisense oligonucleotides (ASOs) have been used. These are single-stranded molecules that act at RNA level by specifically correcting or modifying the expression of the target protein to a less deleterious form^41,42^. ASOs have been therapeutically used to reduce protein production whether via RNase H cleavage or mRNA-ASO complex to block mRNA-ribosome interaction. It is also possible to modulate the splice signalling and among the different types of ASOs, Phosphorodiamidate Morpholino Oligomers (PMOs) have highly desirable molecular properties that make them suitable for this last approach. PMOs are DNA-analogue molecules that comprise a nucleic acid base, a methylenemorpholino ring and phosphodiester inter-subunit bonds are replaced by phosphorodiamidate linkages, all of which produce a structure that provides high-targeting specificity, stability, low toxicity and resistance to nucleases, assuring the maintenance of a good long-term activity within the cell^43^.

In this study we propose an approach to test *in vitro* antisense therapy for NF2. This consists of forcing the skipping of exons harbouring a frameshift or nonsense variant while preserving the gene reading frame, being potentially applicable to 9 of the 15 *NF2* exons in which truncating variants have been reported. Thus, taking into consideration the NF2 genotype-phenotype association, frameshift or nonsense variants that generate a more severe phenotype could be altered at RNA level and potentially generate less deleterious protein forms, resulting in a mitigation of phenotype severity. In addition, we tested the feasibility of modulating and correcting the aberrant splicing of the *NF2* gene originating from different point-mutations located near canonical splice sites.

## RESULTS

### Antisense therapy strategies for *NF2*

The first approach of this study aimed to test if PMOs could be used to specifically force the skipping of in-frame exons carrying a truncating variant, thus maintaining the *NF2* gene reading frame, and test if the generated Merlin protein could act as an hypomorphic form, still preserving partial function and to some extent, able to rescue an *in vitro* phenotype. In this way, taking into account that nonsense or frameshift variants are associated to more severe phenotypes than in-frame variants, based on genotype-phenotype correlations, these truncating variants could be modified to less deleterious forms (Figure 1A), with the need to assess whether the effect of the exon skipping on both alleles was more beneficial than harbouring a truncating variant in heterozygosis.

**Figure 1.**
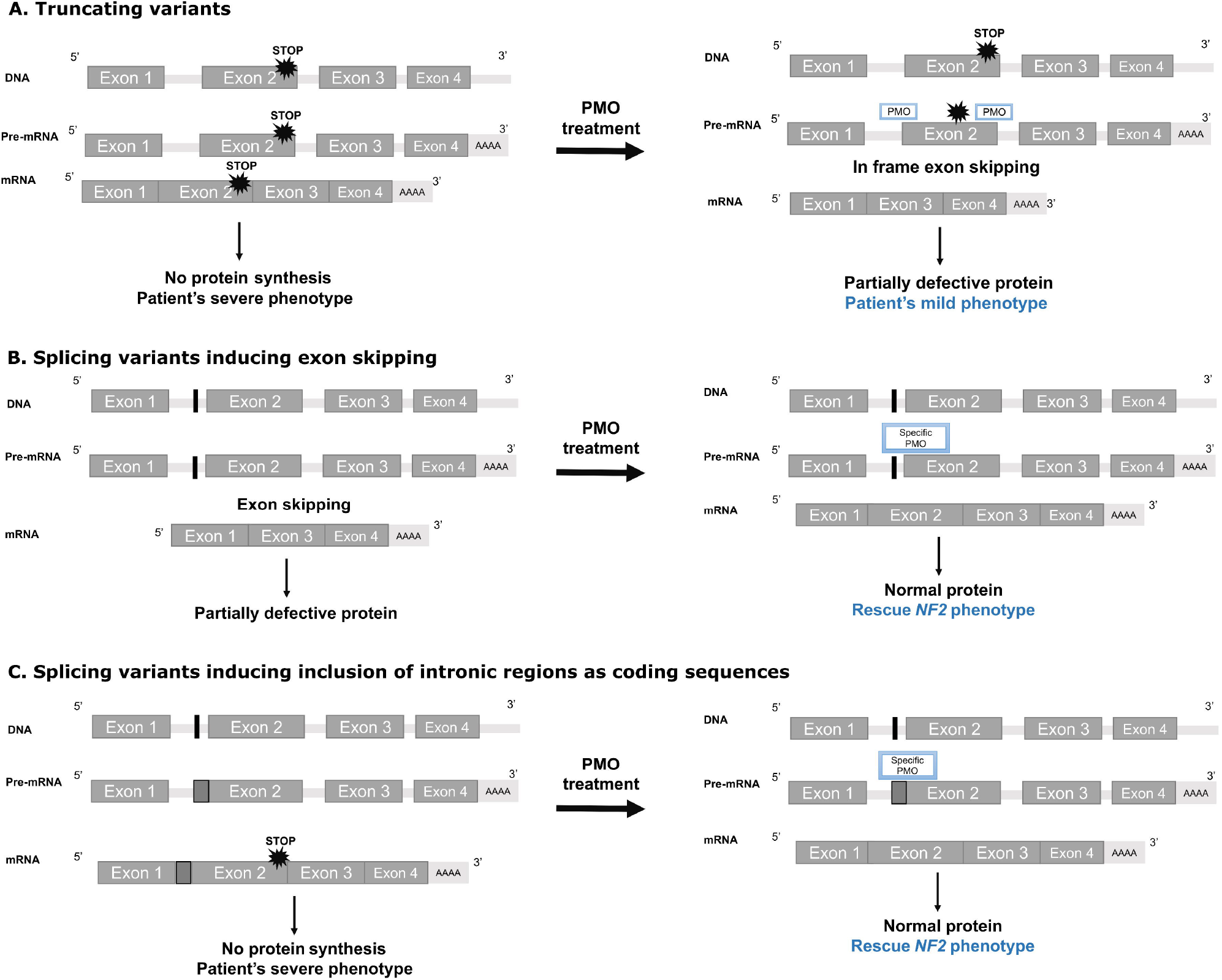
Experimental approaches to test antisense therapy for *NF2* variants. (**A) Truncating variants**. In blue, expected outcome after PMO treatment. PMOs are designed at the donor and acceptor sites of an in-frame exon to induce skipping of exons harbouring truncating variants. The generated in-frame deletion is expected to ameliorate the phenotype *in vitro*. (B) **Experimental approach to test antisense therapy for splicing *NF2* variants** that induce exon skipping and result in a truncating protein and (C) that **induce the inclusion of intronic regions as coding sequences**, resulting in alterations in the gene reading frame that finally generate a truncating protein. In blue, expected outcome after PMO treatment. In both approaches, the PMO is expected to mask the aberrant splicing signal and rescue the NF2 phenotype *in vitro*.

A second strategy was evaluated for *NF2* splicing variants. This experimental approach consisted of designing specific PMOs for a particular splicing variant in order to mask the aberrant splice signalling triggered by the pathogenic variant and promote the spliceosome to properly recognize the correct splice site without altering the wild type (WT) allele. This could be used to target splicing variants that induce the skipping of an exon (Figure 1B) or for those that induce inclusion of intronic regions as coding sequences, as previously tested^44^ (Figure 1C).

### Evaluating the use of PMOs for truncating variants affecting the *NF2* gene

We first investigated *NF2* exons that could be appropriate candidates for generating an hypomorphic Merlin isoform after forcing its skipping, while preserving the gene reading frame. The evaluation of the *NF2* transcript (NM_00268.3) identified 9 out of 15 exons that could be skipped while maintaining the translational reading frame. In addition, for the remaining out-of-frame exons it would be possible to induce skipping of two consecutive exons (6&7, 12&13 and 14&15) (Figure S1). All these twelve combinations were prioritized to be *in silico* analysed. We also assessed exon length, since longer exons encode for a more significant proportion of the protein and thus, could cause greater impact on its structure and stability and, as a consequence, functionality. Most *NF2* exons were in the range of normality without any exon overly long (>500bp). We indicated all unique nonsense and frameshift variants described in the publicly available dataset LOVD (Leiden Open Variation Database) and also all truncating and splicing variants identified in our cohort. As previously described, no hotspots were detected and pathogenic variants were spread over the whole gene^16,17^, although several were described in the FERM domain^16,20^ in LOVD database. Next, we annotated Merlin protein post-translational modifications (PMTs) and the theoretical effect of skipping for each exon (or consecutive exons) in the resulting Merlin protein by *PredictProtein*^45,46^. Predicted features are shown in Figure S1. Seven candidate exons showed a high conservation score (>6.5), three were predicted to induce a high change of surface area, while five out twelve exons or combination of them showed several phosphorylation sites, which could be relevant for Merlin regulation. Considering all the different *in silico* indicators, exons 5 and 11 might be the most suitable candidates since they showed less conservation score, did not present any phosphorylation site and the predicted impact over the protein structure would be relatively low. Nonetheless, besides pairs of exons 12&13 and 14&15, the rest of the in-frame exons presented *in silico* metrics relatively similar among them, and therefore, there was no strong evidence for discard any of the exons for further *in vitro* analysis. In the light of this, we tested the effect on Merlin after forcing exon-skipping on all the available patient’s primary fibroblasts with a truncating variant, corresponding to exons 4, 8 and 11.

A pair of PMOs were designed to be complementary to both 5’- and 3’-intron-exon boundaries to induce skipping of exons 4, 8 or 11 of the *NF2* gene (Exon-Specific PMOs) and the induction of the exon skipping was first assessed in fibroblasts from healthy donors (NF2^(+/+)^) (Table 1). These were delivered to primary fibroblasts from healthy donors (NF2^(+/+)^) through an endocytosis-mediated process with the use of the Endo-Porter. The effect of the treatment was considered in relation to the cells treated with Endo-Porter to determine exclusively the effect of the PMO treatment. Dose response and time course studies confirmed the efficiency of the PMO treatment at RNA level, showing full skipping of exons 4 and 8 at 20μM at 24h, 48h and 72h (Figure 2A and Figure S2). For exon 11, two different designs were tested (Table 1; PMO_ES11_v1 and PMO_ES11_v2): while the first (PMO_ES11_v1) failed to induce skipping (data not shown), for the second design (PMO_ES11_v2) the maximum skipping effect observed was at the highest tested dose (40μM), achieving more than 50% of the exon-less form after 72h of treatment (Figure 2A and Figure S2).

**Table 1.**
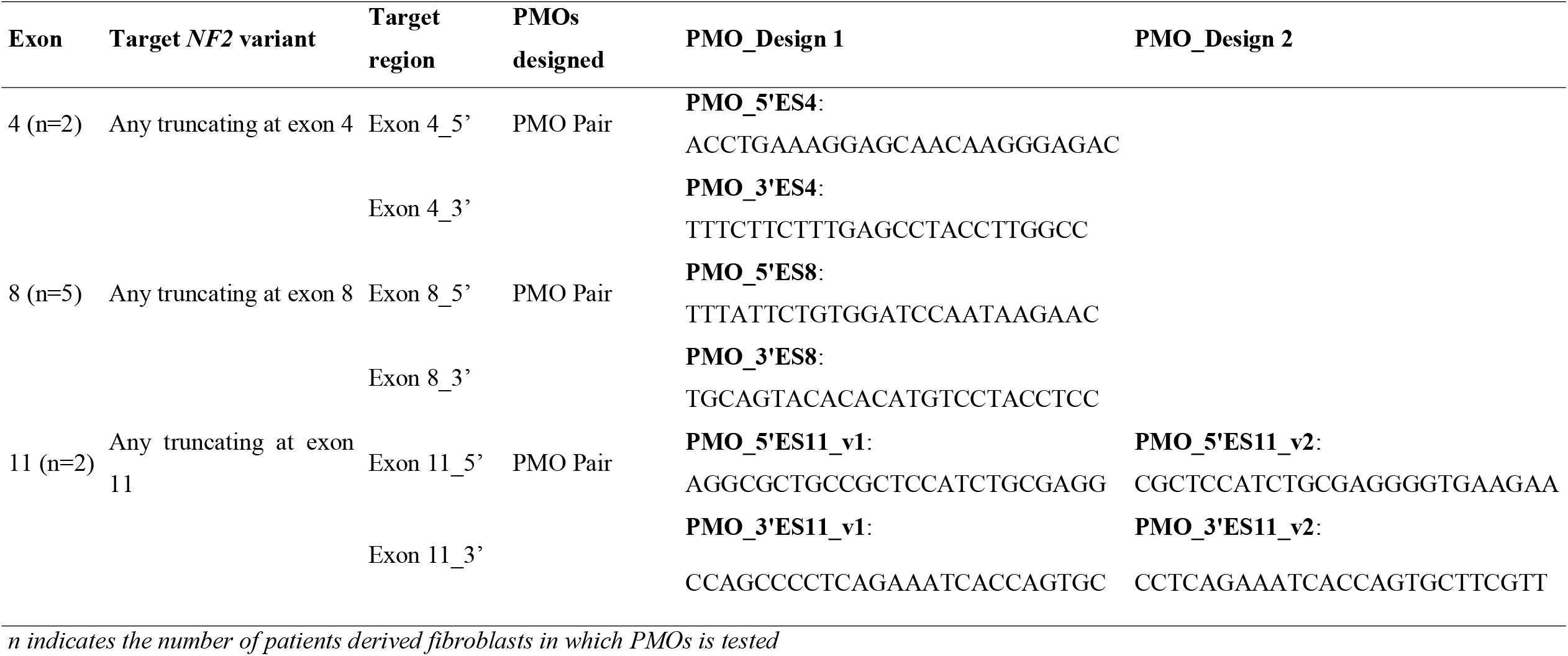
Exon-specific PMOs design to target *NF2* truncating variants.

**Figure 2.**
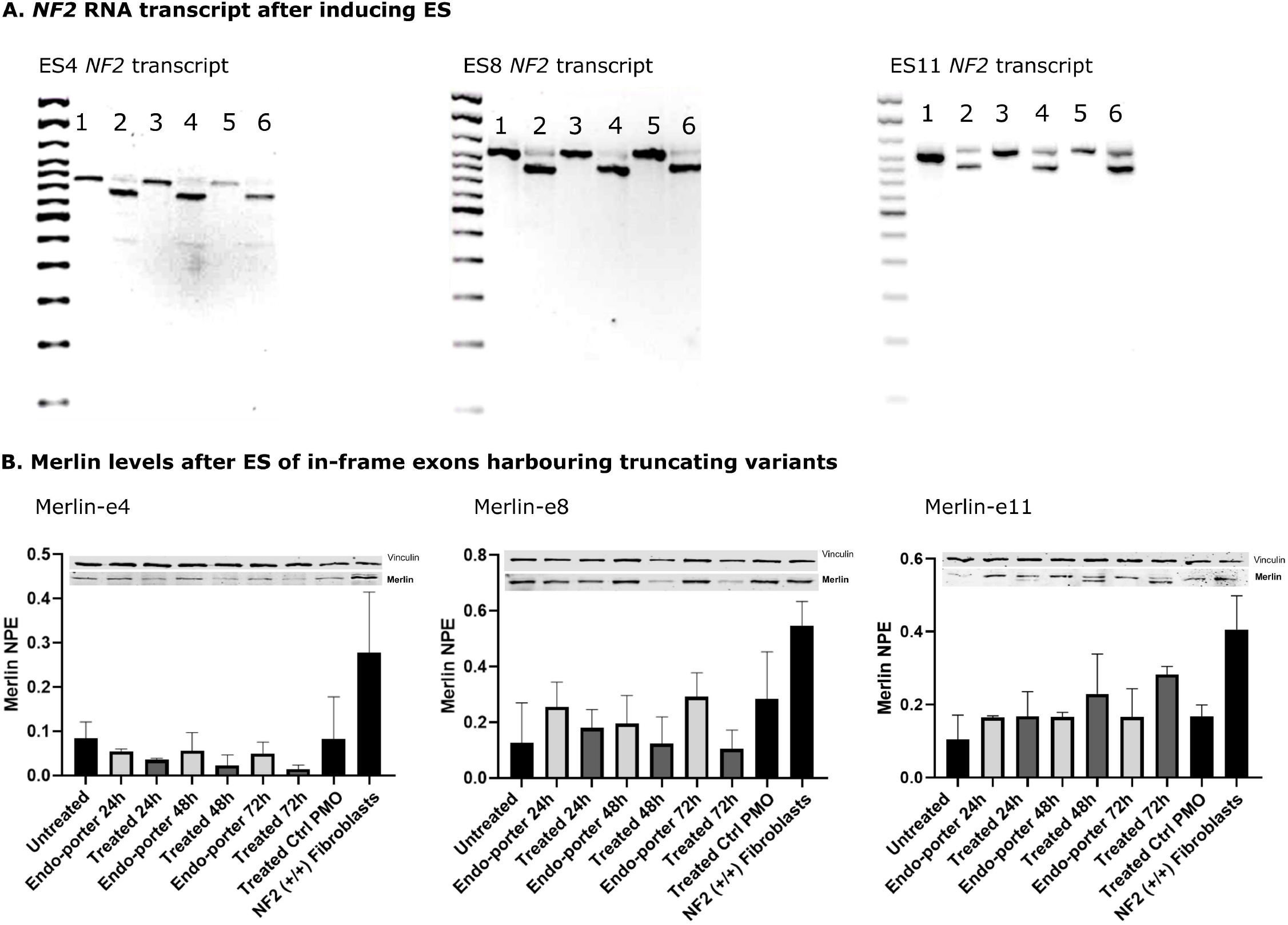
*NF2* RNA transcript and Merlin levels after inducing skipping of in-frame exons harbouring truncating variants. (A) *NF2* RNA transcripts after inducing skipping of exons 4, 8 and 11 of the *NF2* gene showed that we were able to generate these exon-less forms at RNA level. The different lanes correspond to the following conditions: (1) Untreated 24h; (2) Treated (20μM for ES4 and ES8, 40 μM for ES11) 24h; (3) Untreated 48h; (4) Treated 48h; (5) Untreated 72h; (6) Treated 72h. (B) Merlin Western Blot after exon-specific PMOs treatment showed a decrease in Merlin total levels and no presence of exon-less, shorter forms of Merlin-e4 and Merlin-e8. Merlin-e11 was detected and a slight increase was determined in total Merlin levels. Graphs show total protein quantification from three independent experiments in fibroblasts from one patient. Increases or decreases in Merlin levels are considered in respect to the untreated sample. *NF2*(^(+/+)^) fibroblasts stand for control fibroblasts from healthy donors. When indicated “untreated” stands for patient’s fibroblasts (NF2^(+/-)^) without PMO treatment, “Endo-porter” for patient’s fibroblasts (NF2^(+/-)^) treated with PMO’s vehicle, “treated” for patient’s fibroblasts (NF2^(+/-)^) treated with a pair of PMOs for each exon, and “PMO Ctrl” for patient’s fibroblasts (NF2^(+/-)^) treated with a control PMO that does not interfere with *NF2*. ES stands for Exon Skipping and NPE for Normalized Protein Expression.

Next, we analysed whether the skipping of both the *NF2* alleles, the one harbouring a truncating variant and the WT allele, was less deleterious than harbouring a non-functional allele. First, we studied the capacity of restoring Merlin levels *in vitro* after the PMO treatment. Merlin western blot showed progressive loss of protein signal when targeting exons 4 (Merlin-e4) or 8 (Merlin-e8), studied at 24h, 48h and 72h time points in two and five patient samples, respectively (Figure 2B). In order to verify that the induced skipping of exons 4 or 8 caused a reduction of Merlin levels rather than a lack of immunoreactivity; the exon-less *NF2* cDNAs were cloned to a N-terminal FLAG® Tag Expression Vector and transfected to HeLa cells. Flag-Merlin Western Blot showed that there was little or no expression of Merlin-e4 or Merlin-e8, indicating that these new Merlin forms could not be synthesized or were degraded shortly after synthesis (Figure S3).

Conversely, skipping of exon 11 generated a shorter isoform of Merlin (Merlin-e11), and quantification of these results determine a slight increase in the total levels of Merlin (WT Merlin and Merlin-e11) (Figure 2B and Figure S4). These results indicated that a potential hypomorphic Merlin-e11 could result from the skipping of this exon.

### E11 PMO treatment of *NF2*^(+/-)^ fibroblasts partially rescued the phenotype *in vitro*

Functionality of Merlin-e11 was studied in primary fibroblasts harbouring a heterozygous truncating variant in exon 11: one from an adult patient (Patient_ES11_1) and another from a paediatric patient (Patient_ES11_2) (Table S1). Given the complexity of determining differences in severity in the phenotype from studying the signalling pathways in which Merlin is involved [29], physiological read outs were assessed in patients derived fibroblasts (*NF2*^(+/-)^). Due to the role of Merlin in cytoskeletal organization [38,39], Merlin-e11’s functional activity was first tested through actin immunofluorescence. Phalloidin staining revealed an altered phenotype in both primary fibroblast samples, but it was slightly different in each patient tested: patient_ES11_1 showed prominent membrane ruffles, while patient_ES11_2 showed alterations in cytoskeletal abnormalities and a tendency to growth as aggregates, when compared to primary *NF2*^((+/+))^ fibroblasts (Figure 3), and quite similar to the phenotype observed in other primary *NF2*^(+/-)^ fibroblasts (Figure S5). After PMO treatment an improvement of the organizational capacity of the cytoskeleton was observed, membrane ruffles were decreased in patient_ES11_1, and in patient_ES11_2 we observed an improvement in the cell-cell contact organization (Figure 3). Specifically, paediatric derived NF2^(+/-)^ fibroblasts recovered part of the cell-cell contact inhibition and recover the capacity of growth as monolayer after the induction of Merlin-e11.

**Figure 3.**
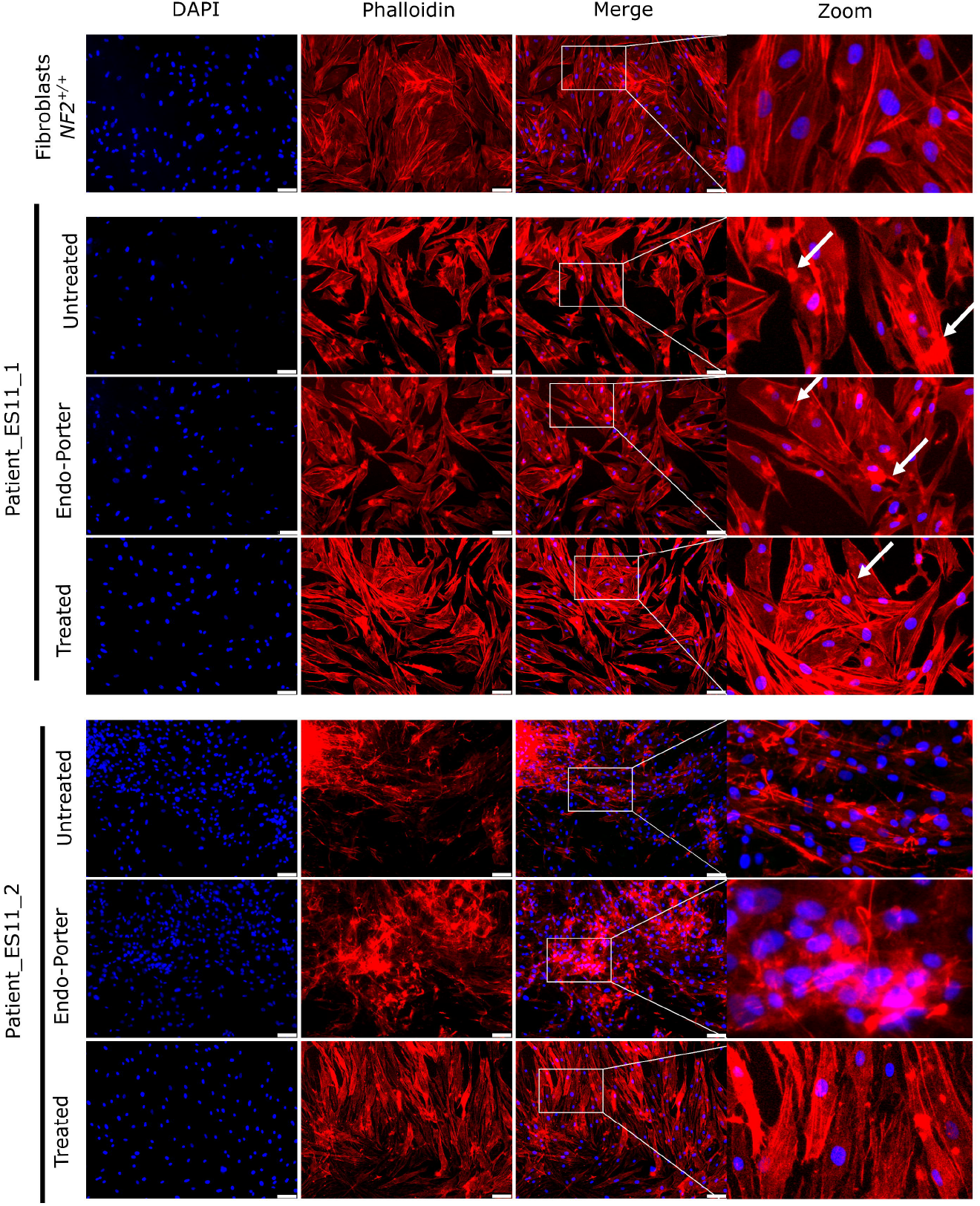
E11 PMO treatment of *NF2*^(+/-)^ fibroblasts induced improvement in actin cytoskeleton organization. Actin immunocytochemistry analysis revealed cytoskeletal abnormalities in patient’s derived fibroblasts. Patient_ES11_1 showed a decrease in membrane ruffle formation and Patient_ES11_2 responded to the treatment showing improved culture organization. DAPI was used to stain cell nuclei and Phalloidin is shown in red. *NF2*(^(+/+)^) fibroblasts stand for control fibroblasts from healthy donors. When indicated “untreated” stands for patient’s fibroblasts (NF2^(+/-)^) without PMO treatment, “Endo-porter” for patient’s fibroblasts (NF2^(+/-)^) treated with PMO’s vehicle and “treated” for patient’s fibroblasts (NF2^(+/-)^) treated with ES11 PMO. Scale bar: 75μm.

In addition, proliferation rates were tested after 72h of treatment by flow cytometry. Patient_ES11_1 showed a statistically significant depletion of proliferation when treated with PMOs (65.6%, p<0.05) (Figure 4A) while patient_ES11_2 showed a less pronounced decrease (25.1%) (Figure 4B). Considering both samples, primary fibroblasts with a truncating variant in exon 11 and expressing Merlin-e11 by PMO treatment showed a 51% reduction in proliferation when compared with the vehicle (Endo-Porter, p<0.01) and 39.7% compared to treatment control (PMO with no effect over the *NF2* gene) (p<0.01), on average (Figure 4C, Figure S6). In order to determine absence of potential toxic effects, a viability test was performed, revealing that PMO treatment did not induce significant differences in cell viability (Figure S7).

**Figure 4.**
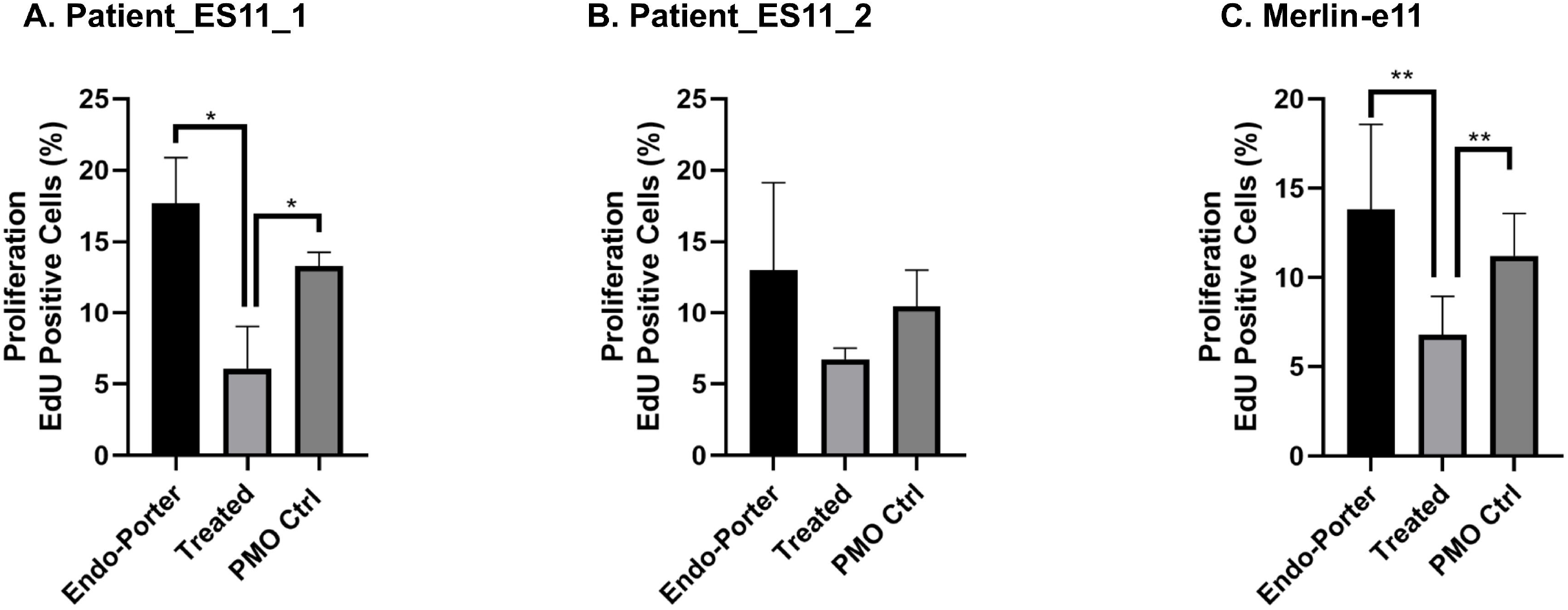
Expression of Merlin-e11 was able to reduce the proliferation capacity of patient fibroblasts. Quantification of EdU-positive cells (percentage over DAPI-positive nuclei). A significant decrease was detected in the proliferation rate of treated fibroblasts when compared to Endo-Porter for Patient_ES11_1 (A). Patient_ES11_2 (B) showed the same tendency. (C) Representation of the mean values of both patients. Mean±SD is represented in the bar, representing three independent experiments. *p<0.05, **p<0.01 denotes statistical significance using unpaired t-test. See also Figure S5. In the figure, “Endo-porter” stands for patient’s fibroblasts (NF2^(+/-)^) treated with PMO’s vehicle, “treated” for patient’s fibroblasts (NF2^(+/-)^) treated with specific PMO and “PMO Ctrl” for patient’s fibroblasts (NF2^(+/-)^) treated with a control PMO that does not interfere with *NF2*.

### Antisense therapy for *NF2* splicing variants

This additional strategy intended to use PMOs, rather than to induce skipping, to mask specifically splicing variants and thus recover the correct splice signal. Two different PMOs were designed for each variant. The first design was a 25-mer variant-specific facilitated by *Gene Tools* according to their guidelines for intron and exon binding subunits and self-complementary moieties (data not shown). We made a second design using splice site predictors to study the variant surrounding sequence (acceptor splicing sites, exonic splicing enhancers (ESEs) or exonic splicing silencers (ESSs)) (based on *Human Splice Finder*) in order to avoid the induction of possible new splicing signals due to the effect of the PMO. In addition, the design avoided masking the canonical splicing GT-AG sites and branch points (YNYYRAY) when possible (Table 2, Figure 5A). We tested the PMOs’ efficacy in primary fibroblasts from NF2 patients harbouring heterozygous germline variants (NF2^(+/-)^) located near the canonical splice site of exons 3, 8 and 15 (Table S1). A dose-response study was performed to test the effect on splicing of different concentrations of PMOs after 24h of treatment, and the efficiency was assessed at cDNA level by RT-PCR. The effect of PMOs on the splicing of *NF2* was analysed by electrophoresis and confirmed by Sanger sequencing, in which the maximum observed effect was at the highest tested dose (20μM) (Figure 5B: Dose Response; Figure S8). A time course was carried out treating fibroblasts with 20μM PMO during 24h, 48h and 72h. PMO treatment in Patient_Spl_1 and Patient_Spl_2 increased the skipping of exons 15 and 8 respectively, in contrast to the desired effect of inducing the inclusion of skipped exon without altering the transcription of the WT allele (Figure 5B: Time Course, B.I and B.II; Figure S8). Similarly, the treatment of PMOs of patients Patient_Spl_3 and Patient_Spl_4, both with nearby variants near the canonical splice site of exon 2, revealed that the variant-specific PMO caused the skipping of both exons 2 and 3 of the *NF2* gene instead of correcting the aberrant *NF2* splicing due to germline variants (Figure 5B: Time Course, B.III and B.IV; Figure S8). Their effect was confirmed by Sanger sequencing (Figure S9). *NF2*^(+/+)^ fibroblasts from healthy donors were also treated with each designed variant-specific PMO and they showed the same effect as *NF2*^(+/-)^ fibroblasts derived from patients (Figure 5B and Figure S8). The effect of the treatment on Merlin was assayed by Western Blot and no improvement in Merlin levels was detected at the studied time points (Figure S10).

**Table 2.**
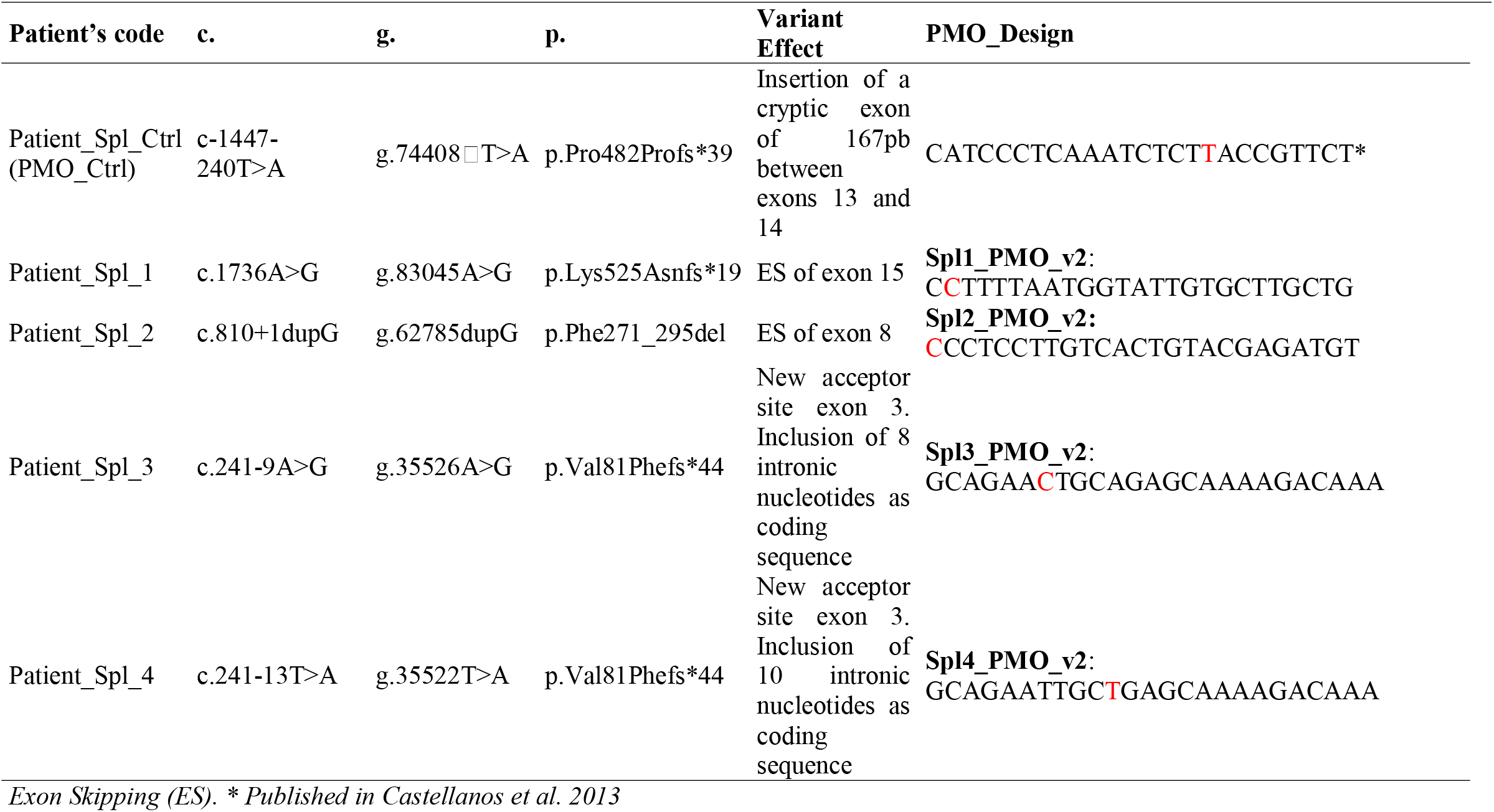
NF2 splicing loss-of-function PMO design.

**Figure 5.**
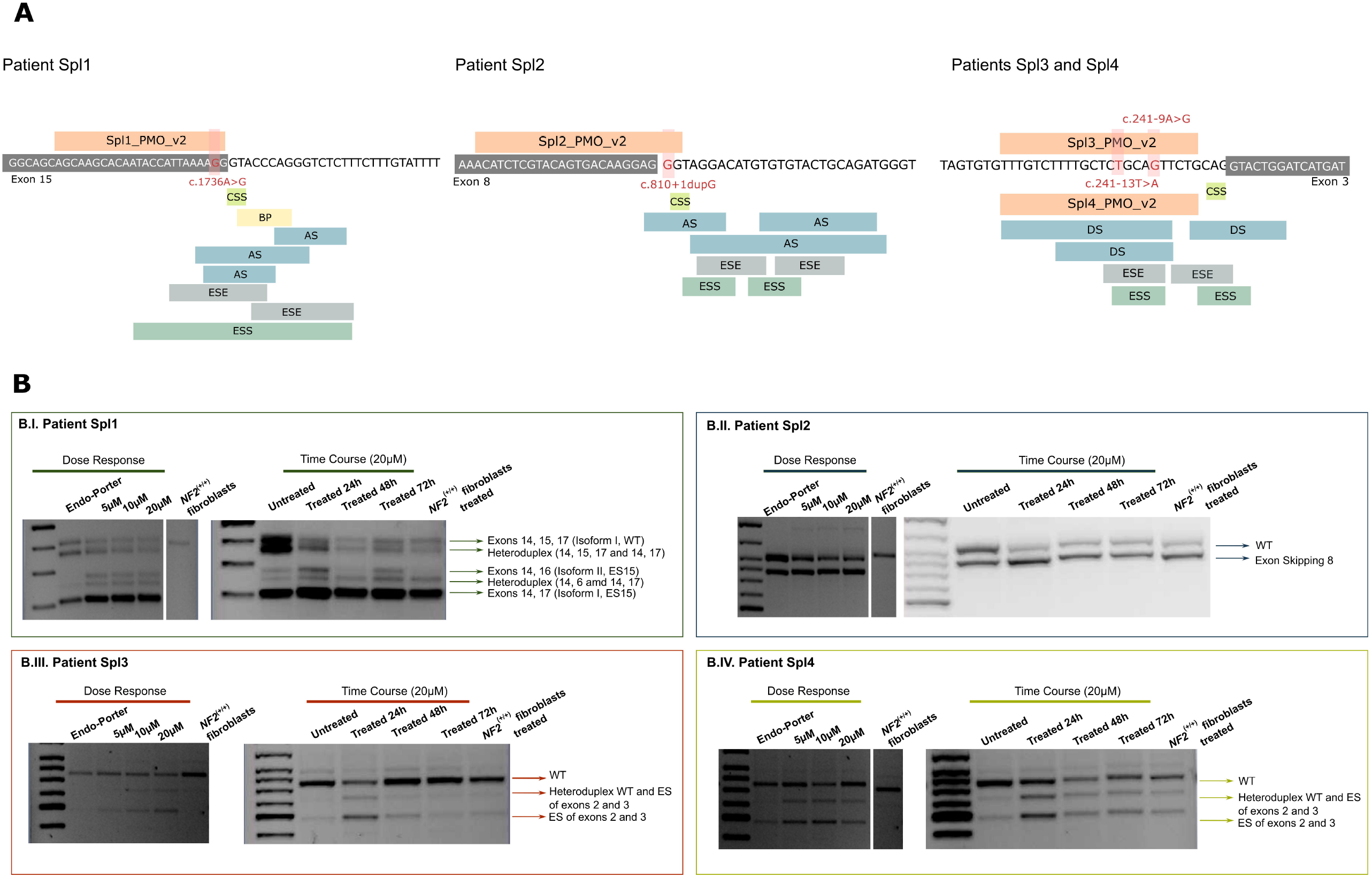
**(A) Design of the PMO specific for each variant.** For the second design (v2), Canonical Splice Sites (CSS), Branch Points (BR), Acceptor and Donor Sites (AS and DS), Exonic Splicing Enhancer sequences (ESE) and Exonic Splicing Silencer (ESS) sequences indicated were taken into account and excluded, when possible, for the sequence design in order to avoid masking the correct splicing signals. The exonic part of the sequence is shown in a grey box and the unlabelled part constitute intronic regions. Tested genetic variants are highlighted in red. **(B) The use of variant-specific PMO did not allow correction of the aberrant splicing signalling**. The effect of the PMOs v2 is shown at cDNA level through dose response and a time course experiments for each patient (Spl_1, Spl_2, Spl_3 and Spl_4). *NF2* transcripts according to the expected molecular weight are indicated next to the bands in the agarose gel. *NF2*(^(+/+)^) fibroblasts stand for control fibroblasts from healthy donors and *NF2*(^(+/+)^) fibroblasts treated indicate that have been treated with the specific PMO of each patient at 20μM. When indicated “untreated” stands for patient’s fibroblasts (NF2^(+/-)^) without PMO treatment, “Endo-porter” for patient’s fibroblasts (NF2^(+/-)^) treated with PMO’s vehicle and “treated” for patient’s fibroblasts (NF2^(+/-)^) treated with specific PMO. ES stands for Exon Skipping and WT for Wild Type.

## DISCUSSION

Neurofibromatosis type 2 (NF2) is a multisystem genetic disorder for which the development of effective therapeutic options with no adverse consequences is needed. The clinical presentation of the disease is variable and related to the type of germline variant inherited in the *NF2* gene^4^. In this context, the development of personalized therapies based on the type of *NF2* variant constitutes an opportunity. In this study, we tested the use of PMOs as antisense therapy to reduce the severity of the effects of *NF2* germline truncating and splicing variants by changing them to milder hypomorphic forms *in vitro* by two different approaches.

In the in-frame exon skipping approach, we targeted nonsense and frameshift variants trying to benefit from the fact that these variants are associated with more severe phenotypes in comparison to in-frame germinal variants in *NF2*^27,29,47^. Thus, we proposed inducing skipping of the exon carrying the truncating variant with the hypothesis of generating potential homozygous hypomorphic forms of Merlin, to ameliorate the pathogenicity of the heterozygous truncating variant *in vitro*. We did not consider mosaic patients in this approach since their phenotype could depend on the type of variant and the percentage of affected cells^32^.

Firstly, we evaluated the theoretical effect of skipping any in-frame exon, and although *in silico* studies did not show any outstanding candidate, the predicted effect on exons 5 and 11 was encouraging. Therefore, we decided to test all exons for which samples were available. The skipping of exons 4 or 8 harbouring *NF2* LOF variants was not able to generate an exon-less Merlin, because Merlin-e4 and Merlin-e8 were degraded. Exons 4 and 8 of the *NF2* gene code for the subdomains B and C respectively of the FERM domain, where the majority of reported pathogenic variants occur^16,20^. The observed results could be due to the fact that the lack of an exon coding for the FERM domain would cause an unbearable aberrant folding of the protein and its synthesis would be prevented or it would be forcibly degraded, although these effects were not predicted *in silico*. In addition, the FERM domain is responsible for the intramolecular interaction with the C-terminal domain, which results in Merlin being in a closed and active conformation^48–50^ to induce the most potential for anti-mitogenic signalling. Thus, if this folding fails, it is possible that the resulting protein would not be able to execute the tumour suppressor activity.

When targeting exon 11 of the *NF2* gene, an exon-less Merlin was expressed (Merlin-e11). Merlin, as a member of the ERM protein family, is involved in the link between actin cytoskeleton and adherent junctions with a relevant role in dynamic cytoskeleton remodelling, such as in membrane ruffling or in the formation of actin micro-spikes^51–53^ Consistent with other studies which described that Merlin loss leads to a dramatic increase of membrane ruffles and actin cytoskeletal disoganization^44,54^, we found that NF2^(+/-)^ primary fibroblasts tested in this study also showed actin cytoskeleton abnormalities. In addition, here we observed a decrease in membrane ruffle formation in response to the expression of Merlin-e11 that could be associated to a partial rescue of the phenotype and an improvement in the cytoskeleton organization after treatment, which could be due to a recovery of the Merlin function as a scaffold protein regulating the linkage between membrane proteins and the cortical cytoskeleton^44,54,55^. Consistent with the role as a tumour suppressor protein, Merlin-deficient cells loose contact inhibition^48,52,56^ and although much remains to be discovered about the adhesive signalling pathways involved, in this study we showed that the induction of Merlin-e11 by PMO treatment contributed to the recovery of cell-cell contact inhibition of proliferation in fibroblasts NF2^(+/-)^ derived from NF2 patients harbouring truncating mutations in exon 11. The tested dose in this study are higher than FDA approved PMOs for other inherited diseases^36,57,58^. Thus, redesigns and modifications of the tested antisense molecule could be performed to improve its efficacy and reduce the required dose, in addition to further pharmacodynamics and pharmacokinetics studies to determine the appropriate range doses and the associated toxicity and effectiveness. Furthermore, treatment effect should be tested in different cell types, especially in schwannoma and meningioma originating cells, as well as *in vivo* studies to determine if the treatment could prevent tumour formation or could reduce tumour burden, when administered systemically before considering the applicability of these molecules in patients

Thus, this study constitutes a proof of concept of a potential medical approach for these variants, although there this is still much to be elucidated about the effect of the PMOs treatment and its therapeutically approach. Further data is required to assess whether this approach could delay the appearance of tumors, cause patients a reduced tumor burden or slower their growth.

A second strategy was directed to splicing variants with the aim of concealing the erroneous splice signal caused by the pathogenic variant and restoring levels of Merlin, in a similar way to the approach that our group used for a *NF2* deep intronic variant^44^ or to induce exon retention by blocking an homozygous single nucleotide variant located 6 nucleotides within exon 7 of *SMN2*^59^. In this study, it has not been possible to design PMOs that permit specifically the correction of aberrant splicing signalling and furthermore, it was not possible to achieve enough specificity to target the mutated allele as the same effect was observed in both patient-derived fibroblasts and control fibroblasts. Based on our results, we hypothesize that when the splicing variant is within +/-13 nucleotides of the acceptor or donor splice site, the PMO could allosterically interfere with the branch point and the U2 snRNA incorporation into the spliceosome^60^. The spatiotemporal localization of the two alleles during splicing could prevent the PMO from specifically targeting the mutated allele and thus it would interfere in the cis-regulation of RNA elements of both alleles equally, resulting in an increased pathogenic effect^61^. Since the approach requires the PMO to be specific for the allele harbouring the pathogenic variant, there is little room for variation in the PMO design for the variants tested in this study.

As far as we know, there are no studies in the literature that have been able to repair the LOF variant effect and induce exon retention through the use of ASOs for a splicing variants located in the canonical splice regions in genes causing autosomal dominant diseases. The most described effect of these molecules targeting intron/exon boundary is exon skipping. It has only been possible through CRISPR/Cas technology by the induction of single nucleotide substitutions^62,63^ Allele-specific approaches have been successfully tested for intronic variants through splice modulation ASOs that mask the erroneous signalling and given their intronic location and distance to splicing regulatory regions, proper protein synthesis can be recovered^44,64–67^. Few works have applied ESSENCE (exon-specific splicing enhancement by small chimeric effectors) method to rescue disease-associated exon skipping and modulate alternative splicing^33,68,69^. Furthermore, for autosomal recessive diseases, such as SMA or DMD^70,71^, or for autosomal dominant diseases with a dominant negative effect, such as Frontotemporal dementia with parkinsonism-17 (FTDP-17) or RP, allele specific ASOs have been used to induce degradation of the disease-causing allele via RNAse H activity and for this purpose, the native DNA backbone is not modified^60,72–74^. Some evidences suggest that Merlin could act in a negative dominant manner through its self-dimerization through FERM-FERM interactions^75–77^, but this seems to be dependent on patient’s variant^78^. In the light of our findings, and if Merlin dominant effect is confirmed, the degradation of the *NF2* mutated allele by the use of RNAse H dependent ASOs could be a possible antisense approach for nonsense, frameshift and splicing pathogenic variants located in the FERM domain, in both in-frame and out-of-frame exons, and then bringing an improvement in the *in vitro* phenotype and opening the door for future gene therapy approaches.

In addition, for those truncating variants that do not exert a dominant-negative effect, the generation of hypomorphic Merlin variants could be beneficial for the patient, and therefore, more variants and hypomorphic Merlin forms should be analysed to understand the contribution of this effect over patient phenotype.

This work is a proof of concept of the use antisense therapy *in vitro* as a personalized therapy for NF2 patients harbouring germline truncating variants at exon 11 of the *NF2* gene. Our approach has shown encouraging results when targeting exon 11, although further characterization assays need to be performed to better characterize the function of Merlin-e11 before considering it for further *in vivo* or pre-clinical assays.

In addition, further studies will be needed to elucidate whether there are other mechanisms or designs to apply ASOs as a therapeutic approach for this disease, as in our experience it has not been possible to recover Merlin levels for variants located at in-frame exons 4 or 8, or to correct splicing signals *in vitro* in the four variants located near canonical splice sites tested, although other splicing variants should be assessed to confirm these results.

## MATERIAL AND METHODS

All procedures performed were in accordance with the ethical standards of the IGTP Institutional Review Board, which approved this study and with the 1964 Helsinki declaration and its later amendments.

### Samples

samples from 14 NF2 patients from the Spanish National Reference Centre (CSUR) on Phakomatoses were included in the study (Table S1) after providing written informed consent. Samples included in the study were those harbouring a splicing variant or a truncating variant in an in-frame exon of NF2.

### Controls

Fibroblasts *NF2*^(+/+)^ stand as non-affected cells for experimental control (WT), with no mutation in the *NF2* gene, obtained from healthy donors.

### Genetic *NF2* test

Genetic testing was performed using the customized I2HCP panel^79^ in blood or tissue when available. To detect splicing variants, patients were studied both at RNA and DNA levels. Variants were validated by Sanger sequencing and analyzed using CLC-workbench (Qiagen, Hilden, Germany) in all primary cultures established using (NM_000268.3; NG_009057.1; LRG_511) as reference sequences.

### *In silico NF2* analysis

NF2 exons were evaluated using NM_000268.3 and NG_009057.1 reference sequences. *NF2* pathogenic variants described in Leiden Open Variation Database [LOVD] in addition with our genetic variants database identified in our NF2 cohort^28^. All *in silico* analysis of exons (and combinations of them) that when skipped, an in-frame Merlin protein is theoretically maintained were based on the analysis performed by Leier et al 2021^80^. Post Translation Modifications (PTM) were retrieved from Phosfosite^81^. Only phosphorylation, acetylation and ubiquitylation were considered as PTMs. Analysis of *in silico* protein modifications by the exon skipping was performed using the web-server version of PredictProtein (https://www.PredictProtein.org)^45,46^.

### Antisense specificity and efficacy

25-mer specific Phosphorodiamidate Morpholino Oligomers (PMOs) for four patients with splicing variants, and a pair of PMOs to induce exon skipping of exons 4, 8, and 11 of the *NF2* gene were designed (NM_000268.3; NG_009057.1; LRG_511), synthesized and purified by Gene Tools (Philomath, OR, USA). Endo-Porter (Gene Tools) was used as vehicle to deliver PMOs into cells at 6μM (manufacturer recommended dose), according to the manufacturer’s instructions. Patient primary fibroblasts carrying *NF2* germline pathogenic variants were treated with the specific single or paired PMOs. A dose-response and a time course study were performed to set-up treatment conditions and the effect of PMOs was assessed by RT-PCR and Sanger sequencing. Levels of Merlin after PMO treatment were analyzed by Western Blot. For all experiments, the correct effect of the treatment at RNA level was confirmed prior to Western Blot analysis.

### Analysis of Merlin-e11 phenotype recovery

an actin immunocytochemistry assay was performed to study the cytoskeletal organization after PMO treatment using Phalloidin (Molecular Probes, Invitrogen). Click-iT™ EdU Alexa Fluor™ 488 Flow Cytometry Assay Kit (Molecular Probes, Invitrogen) was used, according to the manufacturer’s protocol, to determine cell proliferation. Cell viability was assessed using RealTime-Glo™ MT Cell Viability Assay (Promega) following the manufacturer’s instructions.

### *NF2* Cloning and expression

*NF2* gene was cloned using the Gateway® Gene Cloning system (Invitrogen) after treatment to induce the exon-skipping to a FLAG® Tag Expression Vector. The expression vector was transfected into HeLa cells (Lipofectamine LTX with Plus Reagent, Invitrogen) according to the manufacturer’s instructions. Flag expression was assessed by Western Blot (Monoclonal ANTI-FLAG® M2 antibody, Sigma-Aldrich).

### Statistics

Statistical analysis was carried out using GraphPad Prism software v7. The Shapiro–Wilk test was used to test normality and for multiple group comparisons, a two-tailed unpaired t test was performed. Statistical significance is indicated by *p<0.05, **p<0.01, and ***p<0.001. For more information, see Supplementary Materials and Methods.

## ACKNOWLEDGEMENTS

We thank the HGTP Clinical Services and staff for their collaboration in generating and collecting patient samples and data and the Hereditary Cancer Group at the IGTP for their help in improving this work. We thank the IGTP Flow Cytometry core facility and its staff for their contribution and technical support. We would like to acknowledge the constant support of the different NF lay associations: Asociación de Afectados de Neurofibromatosis, Chromo22 and ACNefi. This study has been funded by Instituto de Salud Carlos III through the project PI20/00215 (Co-funded by European Regional Development Fund “A way to make Europe”), Fundació La Marató de TV3 (126/C/2020), the Children’s Tumor Foundation (CTF-2019-05-005), the Spanish Association of NF Affected (AANF), Fundación Proyecto Neurofibromatosis, the Catalan NF Association (AcNeFi), and the Government of Catalonia (2017 SGR 496).

## AUTHOR CONTRIBUTORS

E.C. conceived the study and wrote the manuscript that was revised, corrected, and improved by all the authors. N.C. performed most of the experimental work, analyzed the data, generated the figures for the paper and contributed to writing the manuscript. I.R, S.B., and A.N. performed experimental work. M.T-M. performed bioinformatics analysis. A.P. and H.S. contributed with sample collection. E.S. and I.B. provided scientific input. All authors approved the final version of the manuscript.

## DATA AVAILABILITY STATEMENT

The authors confirm that the data supporting the findings of this study are available within the article [and/or] its supplementary materials.

