## Supplementary Material for "Antisense oligonucleotides targeting exon 11 are able to partially rescue the Neurofibromatosis Type 2 phenotype *in vitro*"

### SUPPLEMENTARY MATERIALS AND METHODS

**Variant analysis:** Human Genome Variation Society (www.hgvs.org) nomenclature guidelines were used to name the mutation at the DNA level, its effect at the mRNA level, and the predicted resulting protein. The first nucleotide of the ATG translation initiation codon is denoted position þ1 according to the NF2 mRNA sequence NM_000268.3 5. and NM_016418.

**Primary cell culture**: For fibroblast isolation, a skin punch was disaggregated into small pieces and digested with 160U/ml collagenase type 1 (Worthington, Lakewood, NJ, USA) and 0.8U/ml dispase grade 1 (Worthington). Fibroblast were grown in Dulbecco's modified Eagle's medium (DMEM; PAA), 10% fetal bovine serum (FBS; Gibco, Paisley, UK), 500U/ml penicillin and 500mg/mL streptomycin antibiotics (Gibco, Paisley, UK) at 37ºC and 5% $CO_2$.

**DNA and RNA preparation and RT-PCR**: Total DNA from peripheral blood was extracted using Flexigene DNA KIT (Qiagen, Hilden, Germany) and DNA from patient-derived cultured fibroblasts was extracted using Maxwell® 16 Cell DNA purification kit according to manufacturer's instructions. For RNA isolation, fibroblasts were treated with 250mg/ml puromycin (Sigma-Aldrich, St Louis, MO, USA) 4–6 h to circumvent the nonsense-mediated decay (NMD) mechanism and extracted using 16 LEV simplyRNA Purification Kit, from Maxwell technology (Promega, Madison, Wisconsin, USA). Total RNA was reverse-transcribed using SuperScript III reverse transcriptase (Invitrogen, Paisley, UK) and random hexamers (Invitrogen).

***In silico NF2* analysis:** NF2 exons were evaluated using NM_000268.3 and NG_009057.1 reference sequences. *NF2* pathogenic variants described in Leiden Open Variation Database [LOVD] v.3.0 Build 27 at May 2022 were considered (https://databases.lovd.nl/shared/variants/NF2/unique), in addition with our genetic variants database identified in our NF2 cohort (most variants have been already described in [1].

All *in silico* analysis of exons (and combinations of them) that when skipped, an in-frame merlin protein is theoretically maintained were based on the analysis performed by Leier et al 2021[2]. Merlin protein original sequence was fetched from https://www.ensembl.org/Homo_sapiens/Transcript/Sequence_Protein?db=core;g=ENSG00000186575;r=22:29603556-29698598;t=ENST00000338641. Post Translation Modifications (PTM) were retrieved from Phosfosite (https://www.phosphosite.org/proteinAction.action?id=1093&showAllSites=true)[3]. Only phosphorylation, acetylation and ubiquitylation were considered as PTMs. Exons with phosphorylation sites are highlighted in yellow on Figure S1. Analysis of *in silico* protein modifications by the exon skipping was performed using the web-server version of *PredictProtein* (https://www.PredictProtein.org)[4,5]. Predicted secondary structure represents the percentage of aminoacid in the exon-skipped remaining merlin protein, that are predicted by *PredictProtein* a change in the secondary structure when compared with the full-length human merlin: *predicted solvent accessibility (%)* accounts for the percentage of aminoacids that changes the solvent accessibility compared to the full-length merlin; *surface contributed by exon* (Å2) predicted solvent accessibility in squared angstroms attributed to the amino acids that have been translated from the exon(s); *change of surface area (Å2)* is calculated by subtracting both the predicted solvent accessibility of the skipped exon and the total merlin without the skipped exon to the full-length merlin: *predicted* $O \rightarrow D$ metric shows the number of aminoacids of the the exon-skipped merlin (compared to the full length) that goes from ordered to disordered; *predicted* $D \rightarrow O$ shows the opposite trend; *predicted P-P binding sites* refers to the number of predicted protein-protein binding sites for each exon; *average conservation score* (range 1-9, being 1 more conserved and 9 less conserved) represents the mean of the conservation score for each merlin exon individually. This value is calculated by *ConSurf*.

**Antisense molecules design and treatment conditions:** In order to test the PMO treatment, cells were seeded at $3 \cdot 10^5$ cells in a six-well plate. 24h later, culture medium was replaced with fresh medium containing the indicated concentration of PMOs (see Results section) plus 6mM of Endo-Porter. PMO reported in Castellanos et al. 2013 was used as an experimental control (PMO_Ctrl). Each experiment was performed in triplicate.

**Western Blotting**: cells were lysed with RIPA buffer (50 mM Tris-HCl (pH 7.4), 150 mM NaCl, 1mM EDTA, 0.5% Igepal CA-630) supplemented with 3mM DTT (Roche, Mannheim, Germany), 1mM PMSF (Thermofisher, Waltham, Massachusetts, USA), 1mM sodium orthovanadate (Sigma-Aldrich), 5mM NaF (Thermofisher), 10 ug/ml leupeptin (Sigma-Aldrich), 5ug/ml aprotinin (Sigma-Aldrich) and 1xPhosSTOP (Roche).

50 µg of protein extracted from primary cultures was loaded to SDS-PAGE (150V) and transferred onto PVDF membranes (1 hour 350 mA at 4oC). Odyssey Blocking Buffer TBS (LI-COR, Lincoln, Nebraska, USA) was used to block the membranes. Primary antibodies were incubated at 4ºC overnight. Membranes were later incubated with IRDye 680LT and IRDye 800CW secondary antibodies (1:1000 dilution, LI-COR) for 1h at room temperature and scanned and analyzed using the Odyssey Infrared Imaging System (LI-COR).

**Western Blot Primary antibodies:** α-NF2: NF2/Merlin antibody (ab88957) α-mouse (Abcam). α-vinculin: anti-vinculin antibody [EPR8185] (ab129002) – α – rabbit (Abcam) was used to normalize protein expression among samples.

**Immunofluorescence**: cells were fixed in 4% para-formaldehyde in PBS for 15min at RT, permeabilized with 0.1%Triton-X 100 in PBS for 10 min at RT, blocked in 10% FBS in PBS for 15 min at RT and stained Alexa Fluor 594 Phalloidin (Molecular Probes, Invitrogen). Nuclei were stained with DAPI and images captured using LEICA DMIL6000 and LASAF software.

**Cell proliferation**: Cell proliferation was determined using the Click-iT™ EdU Alexa Fluor™ 488 Flow Cytometry Assay Kit (Molecular Probes, Invitrogen) according to the manufacturer's protocol and DNA content was studied with DAPI. $1 \cdot 10^5$ cells were seeded in a six-well plate, treated 24h later with the corresponding concentration of PMO. 48h after treatment, 5uM of EdU was added to the cell culture media and incubated for 24h. After 72h of PMO treatment and 24h of EdU incubation, cells were recollected, proceeding according to the manufacturer's instructions and analyzed by FACS. Data was analyzed using an BD FACSCanto™ and BD FACSDiva 6.2 software.

**Cell viability**: Cell viability was assessed using RealTime-Glo™ MT Cell Viability Assay (Promega) according to the manufacturer's instructions. 500 cells were seeded in a ninety-six well plate, treated 24h later with the corresponding concentration of PMO and analyzed at 24h, 48h and 72h after treatment using Flash Thermo Scientific Varioskan® (Thermofisher).

### SUPPLEMENTARY TABLES AND FIGURES

**Table S1** Samples from NF2 patients included in the study and description of the pathogenic variants.

| Sample_Patient | g. | c. | p. | Affected Exon | FGSS |
|---|---|---|---|---|---|
| Patient_ES4_1 | g.38716C>G | c.432C>G | p.Tyr144X | 4 | 6 |
| Patient_ES4_2 | g.43663_43679dupTAGATGAAAA | c.380_396dupTAGATGAAAA | p.Cys133* | 4 | 6 |
| Patient_ES8_1 | g.57759C>T | c.784C>T | p.Arg262X | 8 | 6 |
| Patient_ES8_2 | g.57759C>T | c.784C>T | p.Arg262X | 8 | 6 |
| Patient_ES8_3 | g.57759C>T | c.784C>T | p.Arg262X | 8 | 6 |
| Patient_ES8_4 | g.57759C>T | c.784C>T | p.Arg262X | 8 | 6 |
| Patient_ES8_5 | g.57759C>T | c.784C>T | p.Arg262X | 8 | 6 |
| Patient_ES11_1 | g.73367_73373del | c.1096_1102del | p.Glu366Glnfs*7 | 11 | 6 |
| Patient_ES11_2 | g.30067836C>T | c.1021C>T | p.Arg341* | 11 | 6 |
| Patient_Spl_1 | g.83045A>G | c.1736A>G | p.Lys525Asnfs*19 | 15 | 4 |
| Patient_Spl_2 | g.62785dupG | c.810+1dupG | p.Phe271_295del | 8 | 5 |
| Patient_Spl_3 | g.35526A>G | c.241-9A>G | p.Val81Phefs*44 | 3 | 5 |
| Patient_Spl_4 | g.35522T>A | c.241-13T>A | p.Val81Phefs*44 | 3 | 5 |
| Patient_Spl_Ctrl (PMO_Ctrl) | *positive control ( g.74408 T>A ) | c-1447-240T>A | p.Pro482Profs*39 | 13 | 4 |

FGSS: Functional Genetic Severity Score

| Domain | FERM | α-Helical | CTD |
|---|---|---|---|

| Exon | 1 | 2 | 3 | 4 | 5 | 6 | 7 | 8 | 9 | 10 | 11 | 12 | 13 | 14 | 15 |
|---|---|---|---|---|---|---|---|---|---|---|---|---|---|---|---|
| Exon lenght (nt) | 114 | 126 | 123 | 84 | 69 | 83 | 76 | 135 | 75 | 144 | 123 | 218 | 106 | 128 | 163 |
| Lab truncating variants* | 1 | 5 | 0 | 5 | 2 | 4 | 1 | 2 | 0 | 3 | 2 | 2 | 2 | 2 | 1 |
| Lab CSS variants* | | | 1 | | 2 | | | 1 | | | | | | | 1 |
| Lab intronic variants altering splicing* | | | 2 | | | | | | | | | | 1 | 1 | |
| Unique truncating LOVD reported variants | 5 | 10 | 2 | 1 | 3 | 4 | 1 | 2 | 0 | 1 | 2 | 2 | 2 | 3 | 3 |
| Total truncating LOVD reported variants | 9 | 15 | 2 | 1 | 4 | 15 | 1 | 3 | 0 | 1 | 2 | 2 | 2 | 3 | 3 |
| PTMs (Phosphosite) | 3 | 2 | 1 | 0 | 2 | 0 | | 3 | 2 | 2 | 1 | 9 | | 5 | |
| Predicted secondary structure (%) | 2,7 | 6,5 | 4,2 | 4,6 | 2,4 | 3,5 | | 4,5 | 2,8 | 2,0 | 3,4 | 6,8 | | 2,8 | |
| Predicted solvent accessibility (%) | 0,8 | 0,6 | 1,7 | 0,4 | 0,7 | 1,6 | | 0,2 | 1,2 | 0,4 | 0,3 | 0,8 | | 0,7 | |
| Surface contributed by exons ($A^2$) | 2498 | 2518 | 2538 | 1754 | 1779 | 2787 | | 3149 | 1573 | 2327 | 2566 | 6627 | | 6249 | |
| Change of surface area ($A^2$) | -547 | -291 | -230 | -170 | -260 | -444 | | -51 | -154 | 48 | -133 | -137 | | -451 | |
| Predicted O → D | 6 | 4 | 7 | 7 | 3 | 10 | | 3 | 3 | 3 | 5 | 5 | | 5 | |
| Predicted D → O | 1 | 14 | 7 | 9 | 7 | 0 | | 11 | 15 | 11 | 3 | 20 | | 8 | |
| Predicted P-P binding sites | 2 | 2 | 0 | 0 | 0 | 1 | | 0 | 0 | 3 | 2 | 1 | | 0 | |
| Average conservation score | 4,71 | 7,52 | 6,65 | 7,82 | 4,82 | 7,45 | | 7,97 | 7,97 | 7,64 | 4,24 | 4,45 | | 4,33 | |

**Figure S1.** *In silico* analysis of exons (and combination of them) that when skipped, an in-frame merlin protein is theoretically maintained. Domains of Merlin are shown, as well as the single in frame exons (green) and exons that when skipped together could also maintain the in-frame transcript (blue). * stands for unique variants and CSS for Canonical Splice Site. LOVD: Leiden Open Variation Database. PTM stand for Post Translation Modifications. Only phosphorylation, acetylation and ubiquitylation were considered as PTMs. Exons with phosphorylation sites are highlighted in yellow. Average conservation score ranges from 1 being more conserved up to 9 as less conserved.

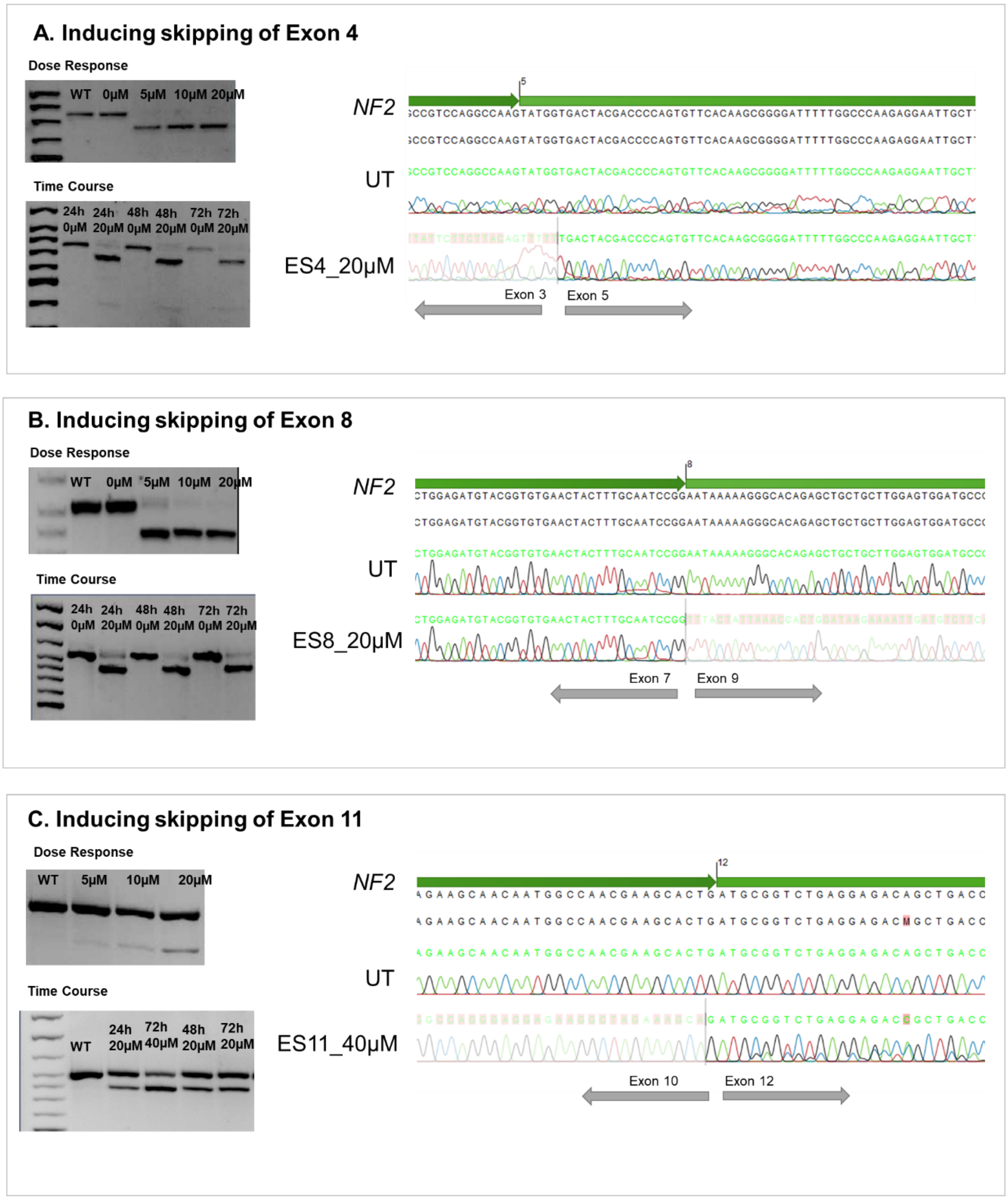


**Figure S2.** Testing the effect of the PMOs to induce exon skipping by a dose-response and a time-course analysis. The bands in the electrophoretic gel reveled a maximum effect of the treatment of exon-skipping 4 and 8 (ES4 and ES8) at 20μM and effect was maintained over 72h at cDNA level (panels A and B). For the skipping of exon 11 (ES11), 40μM of the PMO_5'ES11_v2 and PMO_3'ES11_v2 were used the induce the maximum effect observed that was maintained over 72h (panel C) at cDNA level. Treatment effect was confirmed by Sanger sequencing.

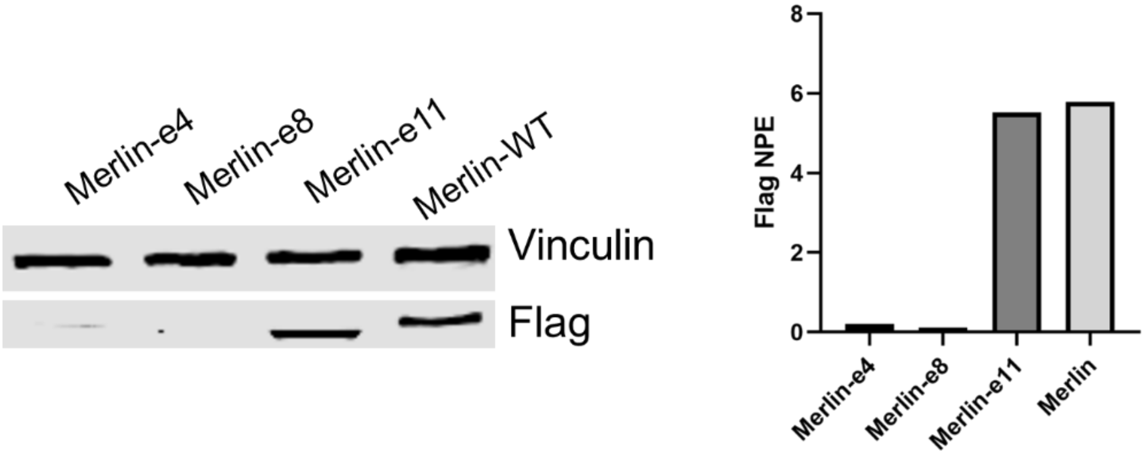


**Figure S3.** Flag-Merlin Western Blot was performed to detect expression of the possible generated hypomorphic forms of Merlin after PMOs treatment. Wild-type Merlin (Merlin-WT) and the potential hypomorphic form of Merlin skipping exon 11 (Merlin-e11) cloned in the N-terminal FLAG® Tag Expression Vector were correctly detected, in contrast to the induced Merlin forms with the skipping exon 4 or exon 8. NPE stands for Normalized Protein Expression.

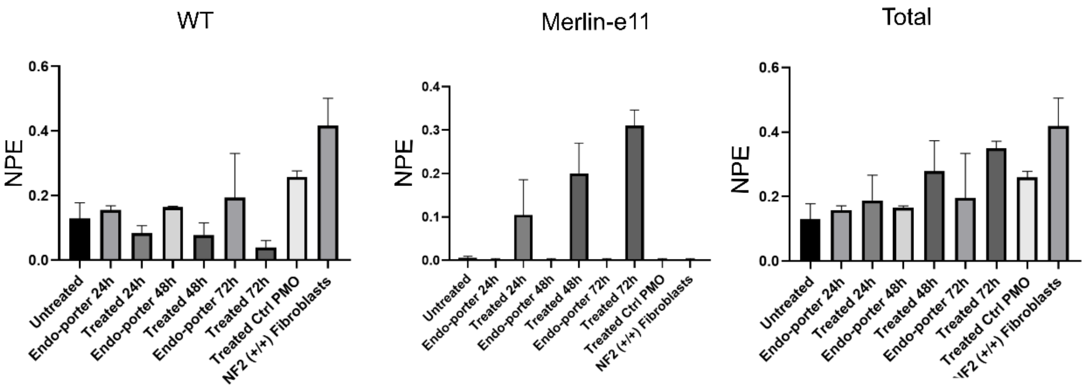


**Figure S4. Quantification of WT Merlin and Merlin-e11** at the different treatment time-points. NPE stands for Normalized Protein Expression and WT stands for wild type. Bars represent the SD from three independent experiments. *NF2*($^{(+/+)}$) fibroblasts stand for control fibroblasts from healthy donors. When indicated "untreated" stands for patient's fibroblasts (NF2$^{(+/-)}$) without PMO treatment, "Endo-porter" for patient's fibroblasts (NF2$^{(+/-)}$) treated with PMO's vehicle, "treated" for patient's fibroblasts (NF2$^{(+/-)}$) treated with ES11 PMO and "PMO Ctrl" for patient's fibroblasts (NF2$^{(+/-)}$) treated with a control PMO that does not interfere with *NF2*. ES stands for Exon Skipping and NPE for Normalized Protein Expression.

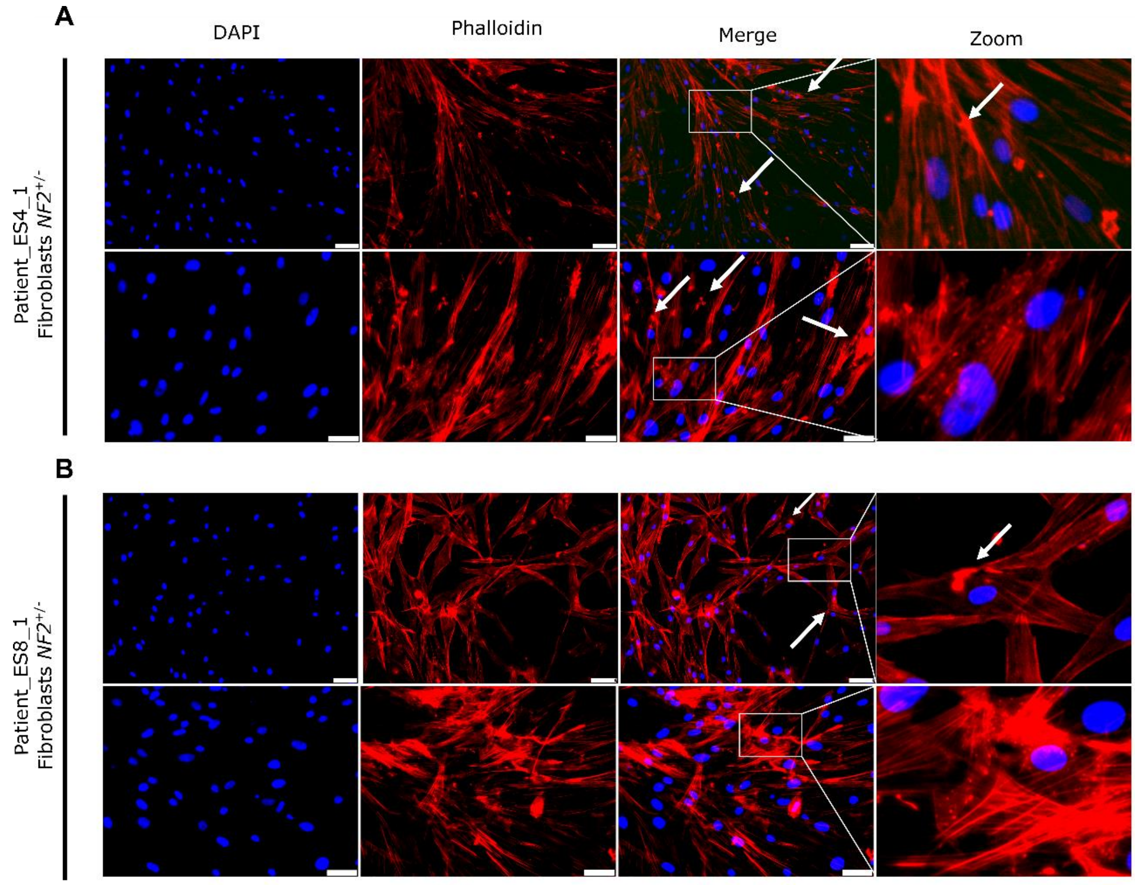


**Figure S5.** Phalloidin staining revealed abnormalities in NF2$^{(+/-)}$ fibroblasts actin cytoskeleton. DAPI was used to stain cell nuclei and Phalloidin is shown in red. Scale bar: 75µm for the upper panel of each patient and 50µm in lower panels. (A) Fibroblasts *NF2*$^{(+/-)}$ from a patient with a truncating variant in exon 4; (B) Fibroblasts *NF2*$^{(+/-)}$ from a patient with a truncating variant in exon 8.

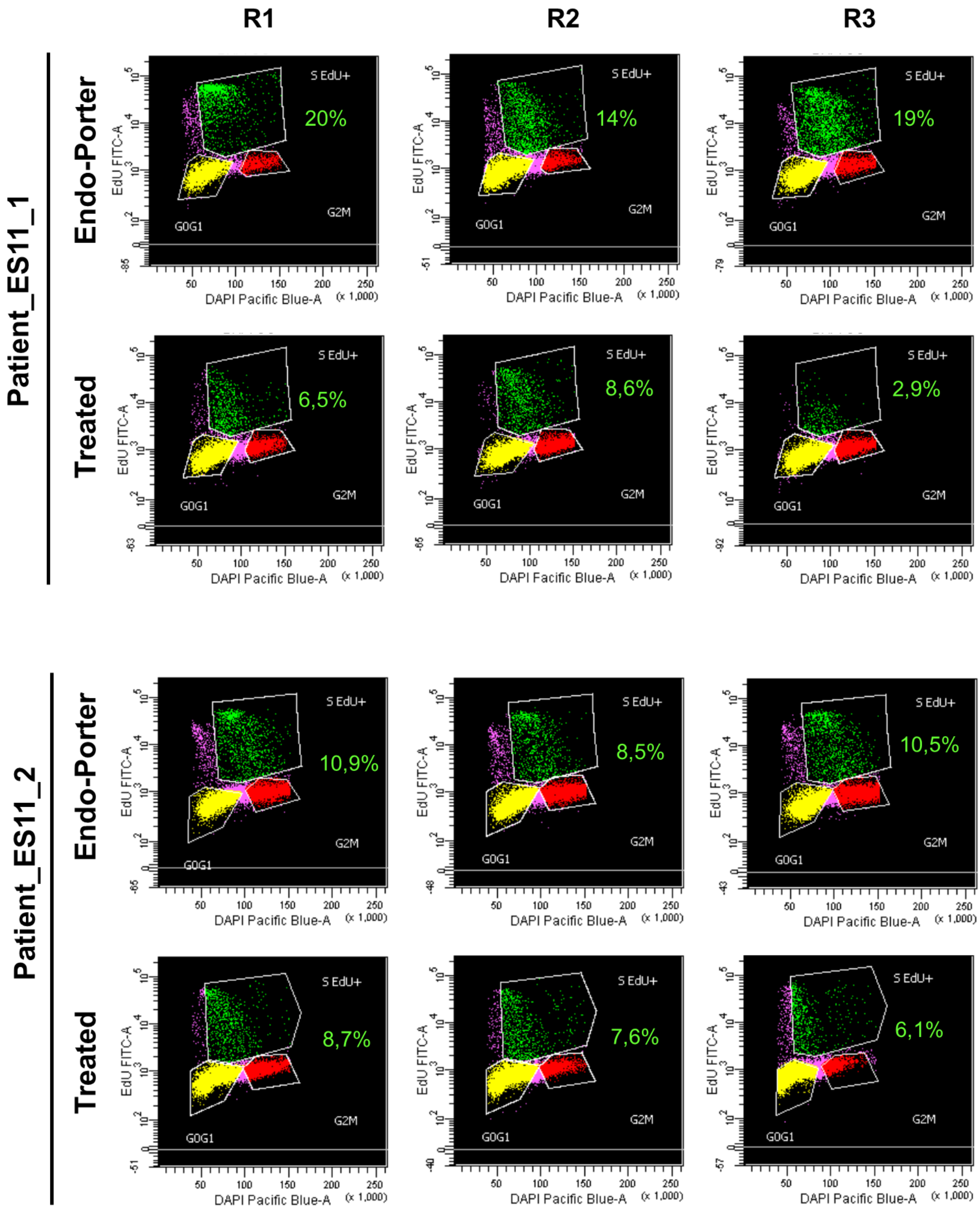


**Figure S6.** Quantification of EdU-positive cells (percentage over total DAPI-positive nuclei) by flow cytometry. Gate were established considering EdU-positive cells and DNA content. Green gate represents EdU-positive cells in S phase, phases G0 and G1 are represented in yellow and red for phases G2 and M. R1, R2 and R3 stand for three independent experiments. When indicated "Endo-porter" for patient's fibroblasts ($NF2^{(+/-)}$) treated with PMO's vehicle and "treated" for patient's fibroblasts ($NF2^{(+/-)}$) treated with ES11 PMO.

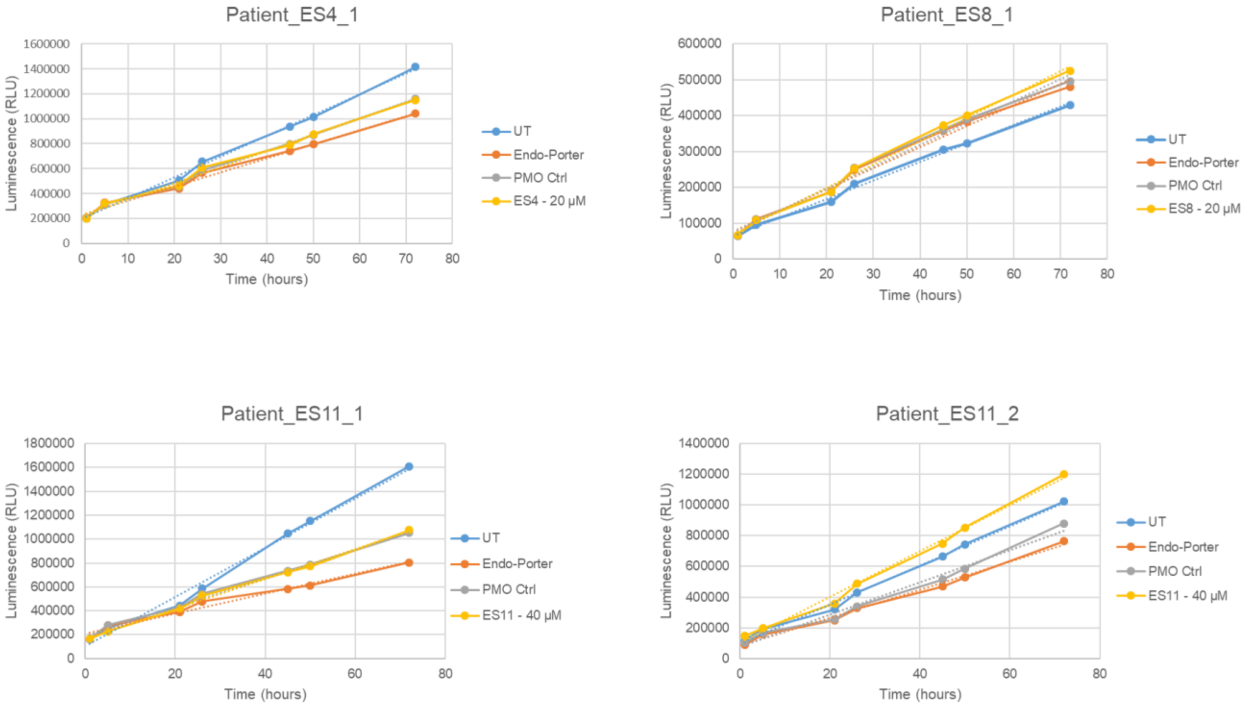


**Figure S7.** Cell viability assay. Results showed no significant differences in cell viability after PMO treatment. The mean of relative luminescence units (RLU) from three independent experiments is represented. Blue stands for patient's fibroblasts ($NF2^{(+/-)}$) without PMO treatment (UT), “Endo-porter” for patient's fibroblasts ($NF2^{(+/-)}$) treated with PMO’s vehicle and is shown in orange, “PMO Ctrl” for patient's fibroblasts ($NF2^{(+/-)}$) treated with a control PMO that does not interfere with *NF2*, shown in grey. Finally,patient's fibroblasts ($NF2^{(+/-)}$) treated with the pair of PMO are indicated as ES4-20µM, ES8-20µM and ES11-40µM and shown in yellow.

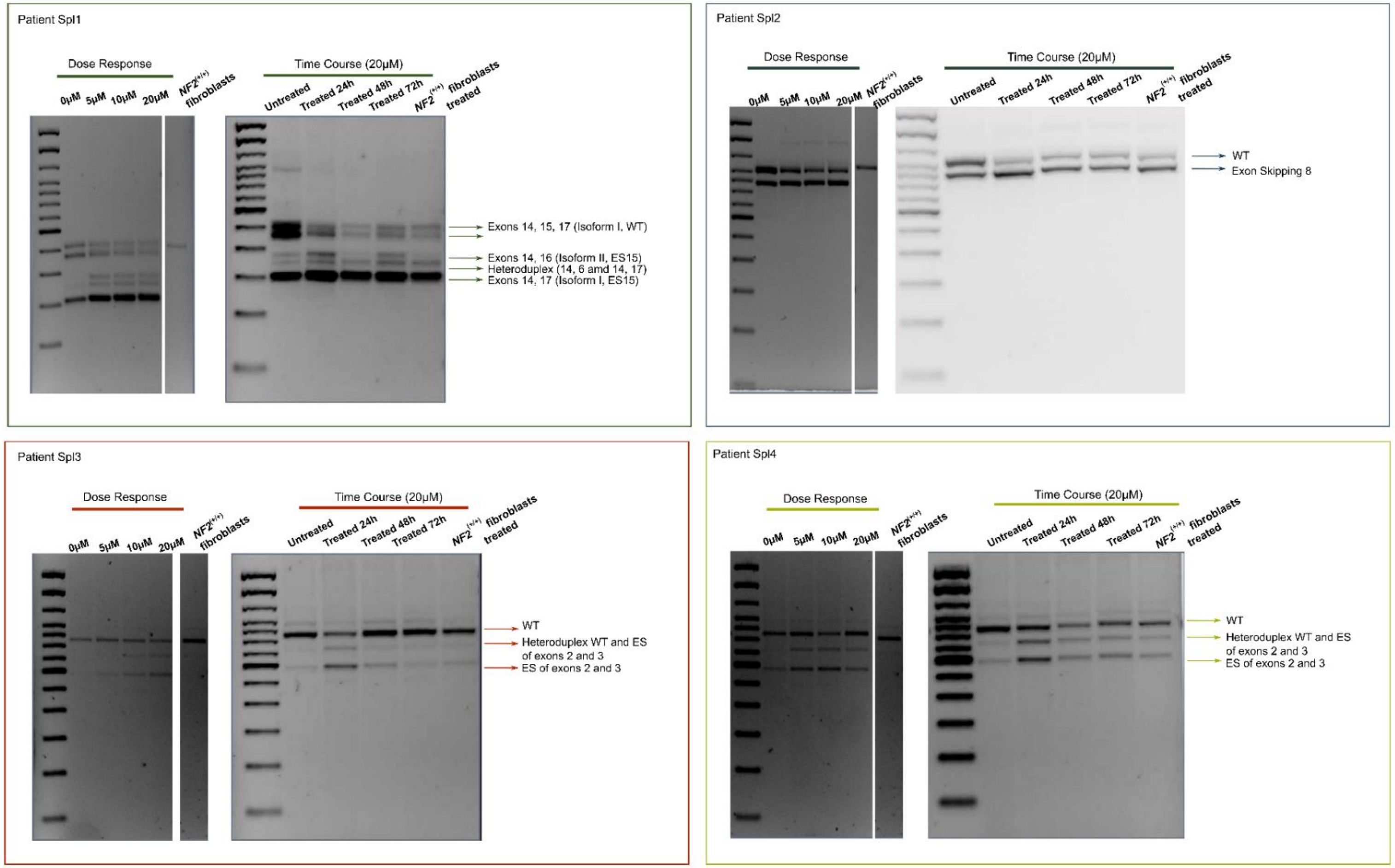


**Figure S8. *NF2* RNA transcripts after PMO treatment in dose response and time-course assays**. The effect of the PMOs v2 is shown at cDNA level through dose response and a time course experiments for each patient (Spl_1, Spl_2, Spl_3 and Spl_4). *NF2* transcripts according to the expected molecular weight are indicated next to the bands in the agarose gel. *NF2*($^{(+/+)}$) fibroblasts stand for control fibroblasts from healthy donors and *NF2*($^{(+/+)}$) fibroblasts treated indicate that have been treated with the specific PMO of each patient at 20µM. When indicated "untreated" stands for patient's fibroblasts (NF2$^{(+/-)}$) without PMO treatment, "Endo-porter" for patient's fibroblasts (NF2$^{(+/-)}$) treated with PMO's vehicle and "treated" for patient's fibroblasts (NF2$^{(+/-)}$) treated with specific PMO. ES stands for Exon Skipping and WT for Wild Type.

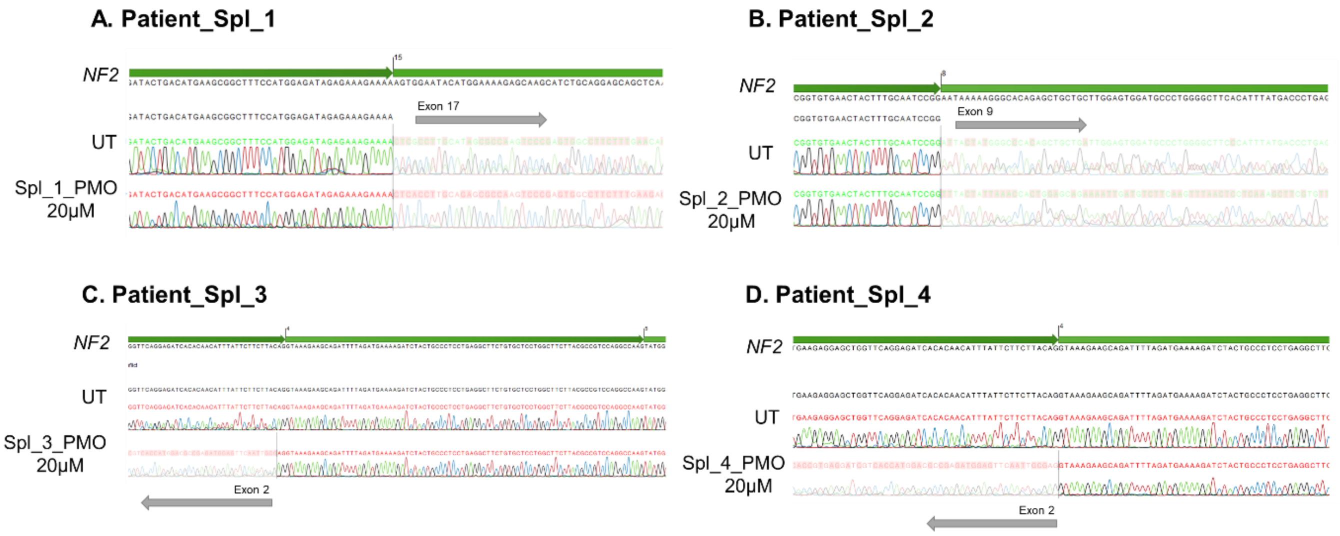


**Figure S9.** Sanger Sequencing confirmed the effect of the variant-specific PMOs observed in the agarose gel.

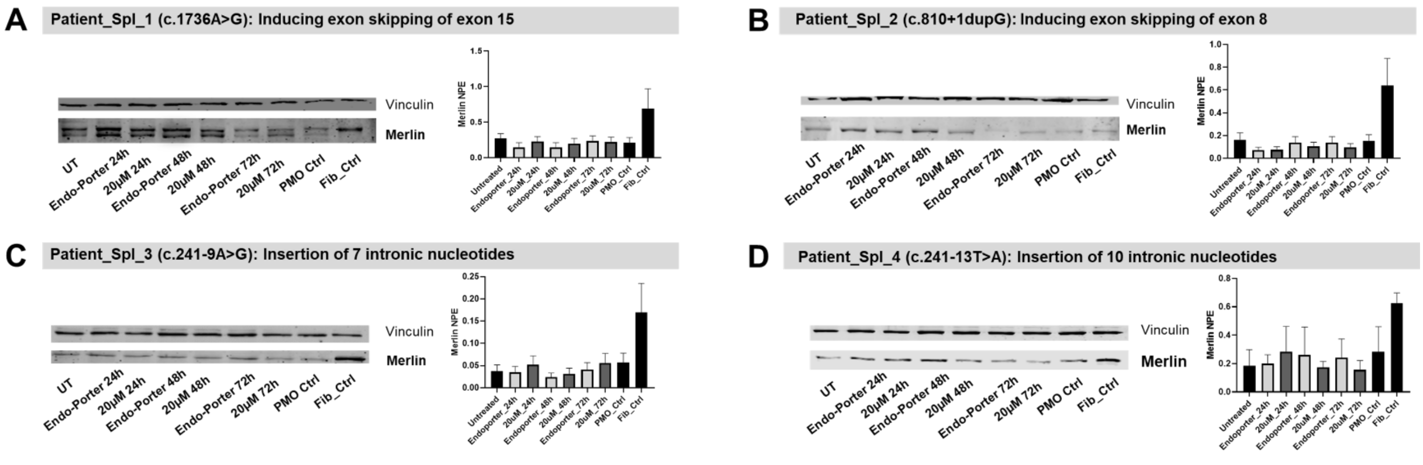


**Figure S10.** Testing variant-specific PMOs targeting NF2 splicing variants. Merlin Western Blot decreased upon PMO treatment. Increases and decreases in levels of Merlin are considered in respect to the untreated sample. Fibroblasts Control (Fib_Ctrl), with no mutation in *NF2* are used as experimental controls. PMO Ctrl stands for patients' fibroblasts treated with a PMO control without any effect in the *NF2* gene. UT stands for Untreated; NPE stands for Normalized Protein Expression. Graphs show protein quantification from three independent experiments.
